# Quantitative intravital imaging of *Plasmodium falciparum* sporozoites: A novel platform to test malaria intervention strategies

**DOI:** 10.1101/716878

**Authors:** Christine S. Hopp, Sachie Kanatani, Nathan K. Archer, Robert J. Miller, Haiyun Liu, Kevin Chiou, Lloyd S. Miller, Photini Sinnis

**Affiliations:** Johns Hopkins Bloomberg School of Public Health, Baltimore, Maryland, USA; Johns Hopkins Malaria Institute, Johns Hopkins Bloomberg School of Public Health, Baltimore, Maryland, USA; Laboratory of Immunogenetics, National Institute of Allergy and Infectious Diseases, National Institutes of Health, Rockville, Maryland; Department of Dermatology, Johns Hopkins University School of Medicine, Baltimore, Maryland, USA; Department of Physics and Astronomy, University of Pennsylvania, Philadelphia, PA, USA; Department of Medicine, Division of Infectious Diseases, Johns Hopkins University School of Medicine, Baltimore, Maryland, USA; Department of Orthopaedic Surgery, Johns Hopkins University School of Medicine, Baltimore, Maryland, USA; Department of Materials Science and Engineering, Johns Hopkins University, Baltimore, Maryland, USA

## Abstract

Malaria infection starts with the injection of motile *Plasmodium* sporozoites into the host’s skin during a mosquito bite. Previous studies using the rodent malaria model indicate that the dermal inoculation site may be where sporozoites are most vulnerable to antibodies, yet, functional *in vivo* assays with human malaria parasites are lacking. Here, we present the first characterization of *P. falciparum* sporozoites in the skin, comparing their motility to two rodent malaria species and investigating whether the environment of its natural host influences *P. falciparum* sporozoite motility using a human skin xenograft model. The combined data suggest that in contrast to the liver and blood stages, the skin is not a species-specific barrier for *Plasmodium*. We observe that *P. falciparum* sporozoites inoculated into mouse skin move with similar speed, displacement and duration, and enter blood vessels in similar numbers as the rodent parasites. Thus, interventions targeting *P. falciparum* sporozoite migration can be tested in the murine dermis. Importantly, to streamline quantification of sporozoite motility, we developed a toolbox allowing for automated detection and tracking of sporozoites in intravital microscopy videos. This establishes a platform to test vaccine candidates, immunization protocols, monoclonal antibodies and drug candidates for their impact on human malaria sporozoites *in vivo*. Screening of intervention strategies for *in vivo* efficacy against *Pf* sporozoites using this new platform will have the potential to validate targets prior to expensive clinical trials.

## Introduction

Malaria, the most deadly parasitic infection of humans, is caused by protozoan parasites of the genus, *Plasmodium*, affecting over 200 million people globally per year (1). The majority of deaths due to malaria are caused by *Plasmodium falciparum*, which is endemic in sub-Saharan Africa. The symptoms of malaria are caused by cyclic multiplication of the parasite in host erythrocytes, yet infection begins, when an infected mosquito inoculates *Plasmodium* sporozoites as it searches for blood (2). Inoculated sporozoites actively migrate through the dermis to enter blood vessels (3), where the circulation carries them to the liver. Sporozoites invade and multiply inside hepatocytes, from which thousands of liver merozoites are released to initiate blood stage infection.

Gliding motility is a striking feature of the invasive stages of Apicomplexan parasites. Of the invasive stages of *Plasmodium*, sporozoites are the most impressive, moving at high speed (1-3 μm/s) for 1 hour, frequently longer. Sporozoite motility is essential for their exit from the dermis and consequently for their infectivity (2–6). Intravital fluorescence microscopy of sporozoites moving through the dermis of laboratory mice is a powerful technique and has permitted qualitative and quantitative studies of the rodent malaria parasite *Plasmodium berghei*: its migration through the skin and its interaction with blood vessels (2–5). These and other studies demonstrate that the dermal inoculation site is where the malaria parasite is extracellular for the longest period of time in the mammalian host, making this a time of vulnerability for the parasite.

Our previous work highlighted significant changes in *P. berghei* sporozoite motility over the first 2 hours after inoculation (3). Upon entering the dermis, sporozoites move with high speed on relatively linear paths, likely to optimize their dispersal. After 20 min in the tissue, sporozoite trajectories become increasingly confined, a motility pattern that likely optimizes contact with blood vessels and blood vessel entry.

Due to emerging drug resistance of *Plasmodium* parasites and insecticide resistance in the mosquito vectors, a highly effective vaccine is widely viewed as a key step towards defeating malaria (7,8). Sporozoite transmission is a significant bottleneck for the parasite, with 10 to 100 parasites being inoculated into the skin and only 20% of these successfully exiting the dermis (8–10), making this early life cycle stage an attractive target. Indeed it has long been appreciated that vaccination with attenuated sporozoites confers protection, a finding that provided the foundation for the development of RTS,S, a subunit vaccine based on the sporozoite’s major surface protein. In Phase III clinical trials RTS,S conferred partial protection against clinical disease and severe malaria (11). Though falling short of community-established goals, this promising first step validated the sporozoite as a vaccine target, with follow-up studies demonstrating that protection was mediated by antibodies targeting the sporozoite’s major surface protein, the circumsporozoite protein (CSP) (12). Previous studies demonstrated that antibodies targeting CSP can immobilize sporozoites *in vitro* and *in vivo* and can exert a large proportion of their protective efficacy in the dermal inoculation site (2,12–14). Though gliding motility is a good target for interventions, it has only been studied in the rodent malaria parasite, and only *P. berghei* has been subject to quantitative intravital imaging (2–5). Here, we present the first *in vivo* motility assessment of *P. falciparum* in mouse skin and grafted human skin in a humanized mouse model and compare the motility of human and rodent malaria sporozoites *in vivo*. Using two rodent malaria species, *P. berghei* and *P. yoelii*, we also describe species-specific differences between the two rodent species. Moreover, we establish a protocol for high-throughput automated analysis of sporozoite motility *in vivo.* These studies establish an *in vivo* platform for the pre-clinical testing of vaccine candidates, monoclonal antibodies and prophylactic drugs targeting the sporozoite stage of the malaria parasite.

## Results

### Imaging *P. yoelii* and *P. falciparum* in the rodent dermis

We first determined whether *P. falciparum* sporozoites are motile in mouse skin and set out to compare their motility to the rodent malaria parasites, *P. yoelii* and *P. berghei*. In order to image *P. yoelii* and *P. falciparum* sporozoites in the dermis, we made new transgenic parasite lines expressing a fluorophore under a strong sporozoite promoter such that sporozoites were sufficiently bright to be visualized by intravital microscopy. We generated a *P. yoelii* line expressing mCherry under control of the *P. berghei csp* promoter (PBANKA_0403200) (Figure S1) and a *P. falciparum* line expressing tdTomato under control of the *P. falciparum peg4* promoter (PF3D7_1016900) (BioRxiv 500793). Intravital microscopy of *P. yoelii* and P*. falciparum* sporozoites in the dermis of mice was performed as previously described for *P. berghei* (3). Figure 1 shows maximum intensity projections of sporozoite motility over 4 min in representative videos started 10 min after sporozoite inoculation (see Videos S1-S3 for the corresponding time-lapse video). Dermal vasculature was visualized by labeling the pan-endothelial junction molecule CD31 by intravenous injection of fluorescently labeled rat anti-CD31 (3,15). As shown, *P. falciparum* sporozoites are motile in mouse skin.

**Figure 1.**
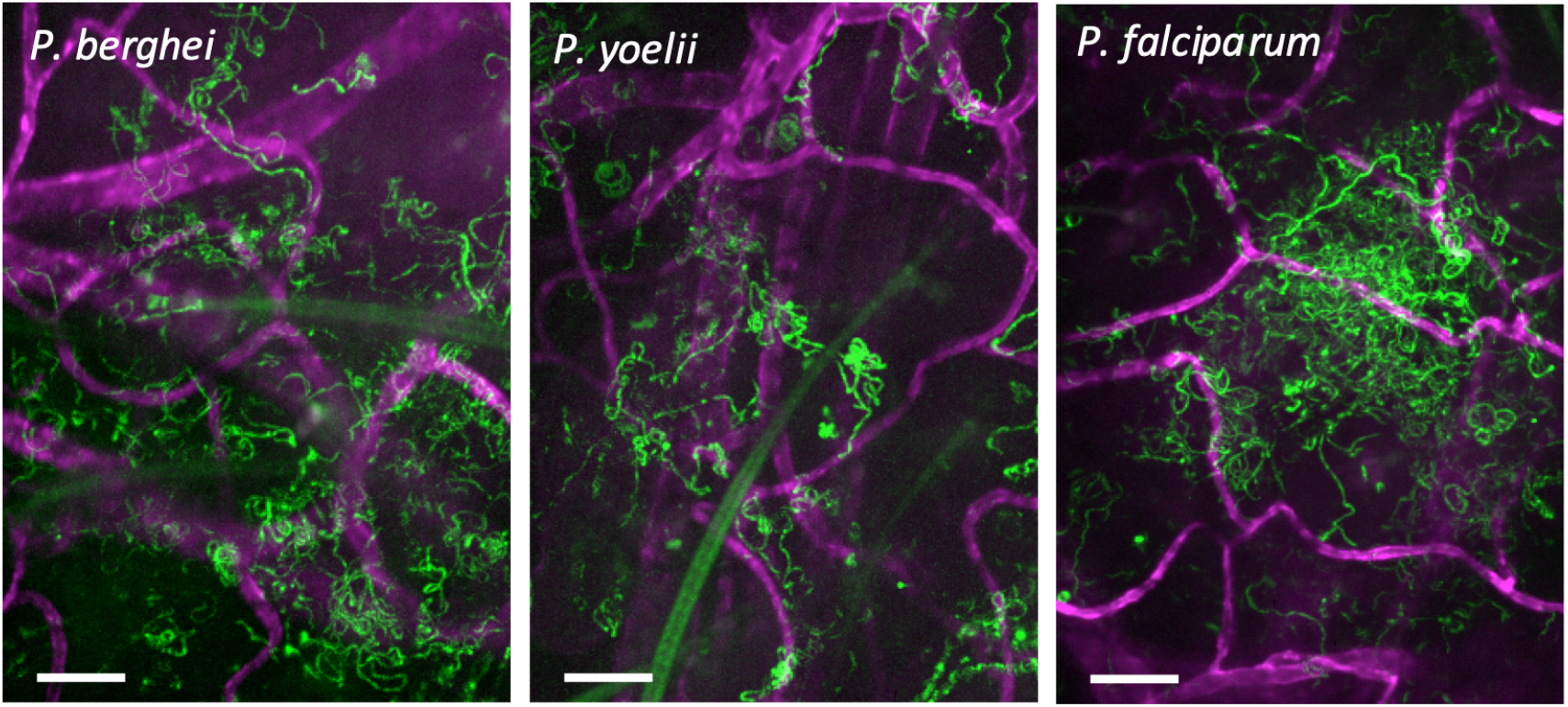
*P. berghei*, *P. yoelii* and *P. falciparum* sporozoites moving in the dermis. Time-lapse microscopy of sporozoites, 10 min after intradermal inoculation into mice with CD31-labeled vascular endothelia. Maximum intensity projection over 240 sec shows trajectories of moving sporozoites (green) and blood vessels (far red). Scale bar, 50 μm. See videos S1-S3.

### Automated sporozoite detection and tracking

To streamline quantification of sporozoite motility, we developed an automated method to detect and track sporozoites. Fiji software was used for vigorous background subtraction and thresholding of raw images, and then spot detection and tracking were performed using ICY software. A visual description of the image processing, spot detection and tracking steps can be seen in Figure S2 and more detailed information can be found in the Supplementary Methods. To verify this new method, we compared the data obtained from the automated spot detection and tracking method to our previous analysis of *P. berghei* motility (3), performed through manual sporozoite detection and tracking, using the same set of videos for both types of analysis. Comparing sporozoite speed and displacement obtained with each method showed that overall the automated method generates data comparable to the data obtained with manual sporozoite tracking (Figure 2). Overall, these data suggest that the automated tracking method can be used to quantify sporozoite motility in a more high-throughput manner than would be possible with time-consuming manual tracking methods.

**Figure 2.**
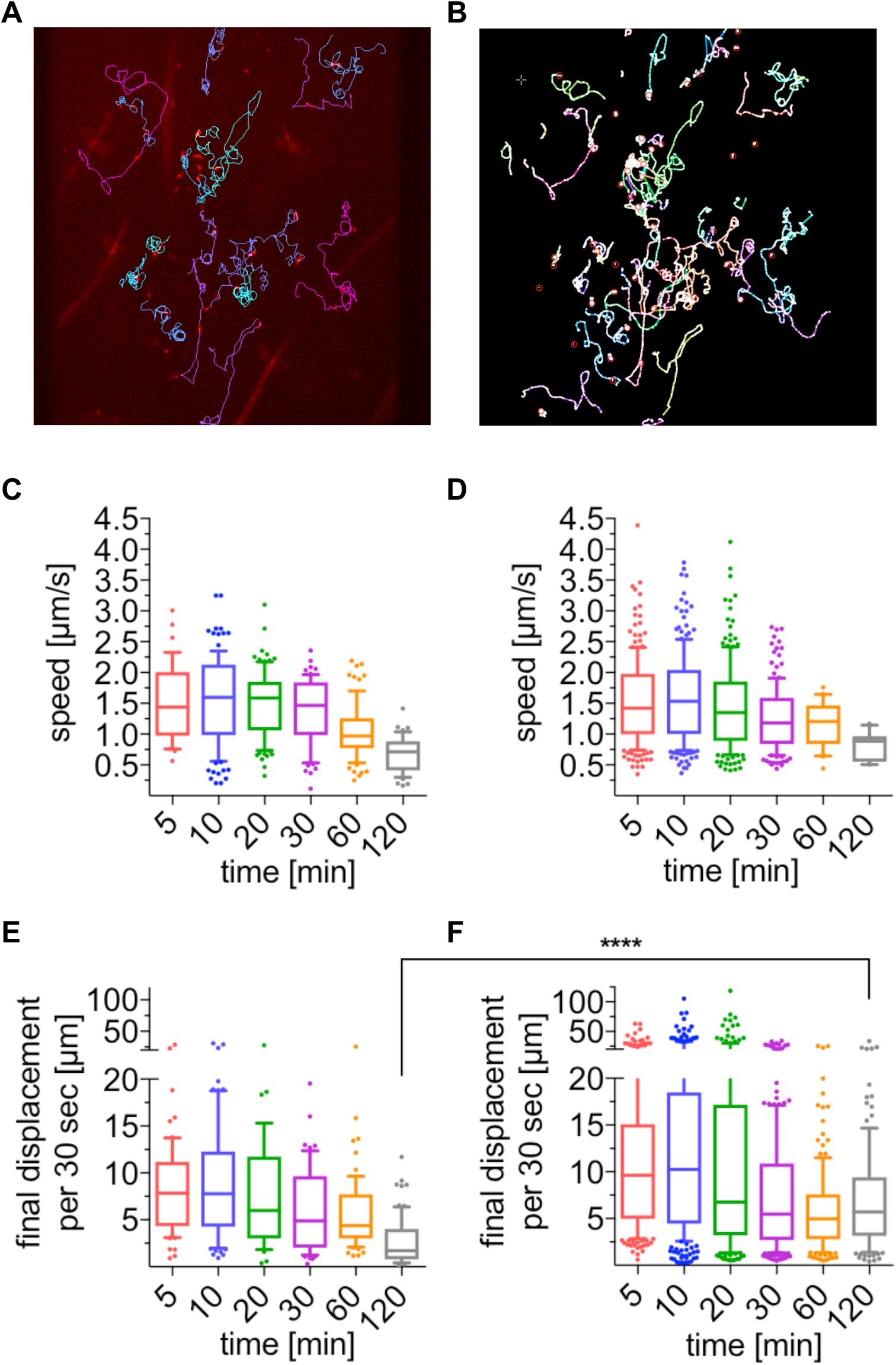
Automatic tracking of *P. berghei* sporozoites moving in the dermis generates data that is comparable to manual tracking. (**A-B**) *P. berghei* sporozoite trajectories obtained with manual (**A**) and automated (**B**) tracking methods. (**C-D**) Apparent speed of sporozoites 5 to 120 min after intradermal inoculation, obtained with manual (**C**) and automatic (**D**) tracking methods. Data is displayed in box and whisker plots with whiskers showing 10 to 90 percentiles and values below and above the whiskers shown individually. Horizontal line indicates the median speed. No statistically significant differences between the corresponding timepoints of manual and automated tracking methods were found (Kruskal-Wallis test). (**E-F**) Displacement of sporozoites 5 to 120 min after intradermal inoculation, obtained with manual (**E**) and automatic (**F**) tracking methods. To normalize between tracks of different duration, displacement is displayed as final displacement per 30 seconds interval. Data is displayed in box and whisker plots with whiskers showing 10 to 90 percentiles and horizontal line showing the median speed. The only statistically significant difference in sporozoite displacement between the manual and automated tracking methods was at 120 minutes (Kruskal-Wallis test, p<0.0001). For panels **C**–**F**, a varying number of videos were processed for each time point: **C**&**D**: 5 min (3 videos/37 manual tracks/203 automated tracks), 10 min (4 videos/112 manual tracks/207 automated tracks), 20 min (4 videos/95 manual tracks/184 automated tracks), 30 min (4 videos/63 manual tracks/155 automated tracks), 60 min (2 videos/77 manual tracks/23 automated tracks), and 120 min (2 videos/44 manual tracks/11 automated tracks). **E**&**F**: 5 min (3 videos/61 manual tracks/203 automated tracks), 10 min (6 videos/74 manual tracks/292 automated tracks), 20 min (4 videos/47 manual tracks/184 automated tracks), 30 min (6 videos/53 manual tracks/235 automated tracks), 60 min (7 videos/65 manual tracks/162 automated tracks), and 120 min (6 videos/48 manual tracks/151 automated tracks).

Comparing sporozoite speed obtained with both tracking methods showed that the parasite speed does not change significantly over the first 30 min and we find a drop in speed at 120 min by both methods (Figure 2C-D). No significant differences in sporozoite speed between corresponding timepoints of the two different tracking methods were found (Kruskal-Wallis comparison) (Figure 2C-D). Nonetheless, some differences in these two data sets are expected, given that the manually-tracked data set included only a subset of the total motile sporozoites that were tracked in the automated analysis: While the automated analysis tracks all motile sporozoites, therefore generating tracks of different total durations, the manual tracking analysis only included motile sporozoites that were observed in the field of view for the full duration of the 4 min long video, which was a necessary requirement to allow calculation of mean square displacement (3). To be able to directly compare sporozoite displacement for tracks of different durations, displacement was normalized to the average displacement per 30-second interval, thus allowing comparison of the automated and manual tracking data sets. As previously described (3), sporozoite displacement drops at 20 min after inoculation in both data sets (Figure 2E-F). However, automated tracking resulted in statistically higher displacements at the latest time point, 120 minutes. This is likely due to inclusion of all motile sporozoites in the automated tracking data set, because sporozoites that leave the field of view would be predicted to have higher displacements than those that stay in the field. To determine if this was the case, sporozoites that were leaving the field of view and thus excluded from the manual analysis, were manually tracked for the 10 min and 120 min time points and added to the previous manual analysis. This showed that the displacement of the total sporozoite population at both 10 min and 120 min after inoculation is higher than was suggested by the original analysis of sporozoites that remain in the field of view throughout the acquired video (Figure S3).

### Constrained motility of *P. berghei*, *P. yoelii* and *P. falciparum* sporozoites over time

Using the automated detection and tracking method, we analyzed motility of *P. yoelii* and *P. falciparum* sporozoites moving in the dermis of mice and compared their motility to the data obtained from automated analysis of our previous imaging data of *P. berghei* sporozoites. Sporozoites moving in the dermis were imaged over the first two hours after intradermal inoculation and videos were acquired at 5 min, 10 min, 20 min, 30 min, 60 min and 120 min after injection (examples shown in Videos S1-S3).

Sporozoite trajectories were re-centered to a common origin to visualize the sporozoite displacement, showing a gradual decrease in displacement over time for all three *Plasmodium* species (Figure 3A-C), suggesting that similar to *P. berghei*, the displacement of *P. yoelii* and *P. falciparum* sporozoites decreases over time after dermal inoculation. While the previous displacement analysis for comparison of the automated to the manual tracking data showed displacement per 30 second interval (Figure 2E&F), this analysis shows the final displacement of motile sporozoites, thus allowing analysis of the total displacement, a parameter that is lost by the normalization to 30 second interval. The analysis of final displacement showed that the median displacement of all three species is comparable, with sporozoites displacing 20 μm, on average, at 5 min after intradermal injection.

**Figure 3.**
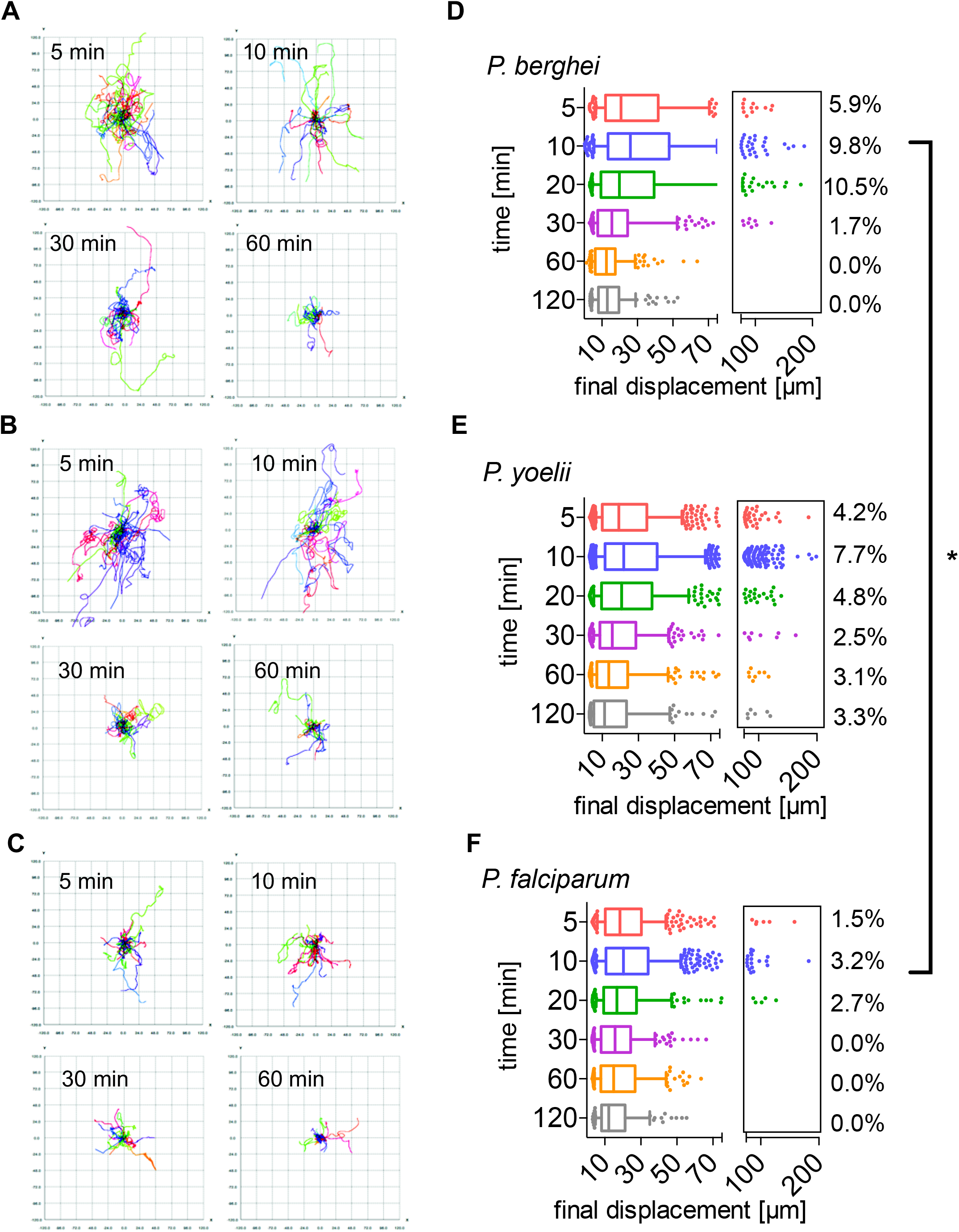
Motility of *P. berghei*, *P. yoelii* and *P. falciparum* sporozoites moving in the dermis is increasingly constrained over time. (**A-C**) Tracks generated through automated tracking were plotted to a common origin, to visualize parasite dispersal *P. berghei* (A), *P. yoelii* (B) and *P. falciparum* (C) at 5, 10, 30 and 60 min after intradermal inoculation. (**D-F**) Displacement of *P. berghei* (D), *P. yoelii* (E) and *P. falciparum* (F) sporozoites 5-120 min after inoculation in box and whisker plots displaying the 10-90 percentile and individual outliers, with horizontal line showing median displacement at each time point. Percentage values in boxes indicate the fraction of tracks displacing over 75 micrometers at each time point. Within a given species, the difference in sporozoite displacement between 5 min and 120 min is statistically significant (Kruskal-Wallis test: *P. berghei* p<0.0001, *P. yoelii* p<0.0001, *P. falciparum p*<0.01). Comparisons between species showed no statistically significant difference in displacements for any given time point, with the exception of the 10 min time point in which there was a significant difference between *P. berghei* and *P. falciparum* (Kruskal-Wallis test, p<0.05). For panels **(D–F)**, a varying number of videos were processed for each time point after inoculation: **(D)** Same dataset as used for Figure 2D: 5 min (3 videos/203 tracks), 10 min (6 videos/292 tracks), 20 min (4 videos/184 tracks), 30 min (6 videos/235 tracks), 60 min (7 videos/162 tracks) and 120 min (6 videos/151 tracks). **(E)** 5 min (7 videos/714 tracks), 10 min (16 videos/1514 tracks), 20 min (4 videos/415 tracks), 30 min (4 videos/319 tracks), 60 min (4 videos/255 tracks), and 120 min (4 videos/180 tracks). **(F)** 5 min (6 videos/389 tracks), 10 min (6 videos/464 tracks), 20 min (3 videos/184 tracks), 30 min (2 videos/195 tracks), 60 min (2 videos/239 tracks), and 120 min (2 videos/103 tracks).

Interestingly, all three species showed the highest final displacements at 10 min after inoculation (Figure 3D-F). After this time point, final displacement gradually drops to 10-15 μm at 120 min after injection. For each of the 3 species, the difference in sporozoite displacements between 5 min versus 120 min, is statistically significant. Comparisons of displacements between species at each time point were not significant with the exception of the 10 min time point, which was significantly different between *P. berghei* and *P. falciparum*.

We also compared the percentage of sporozoites moving with very high displacements, between 75-200 μm (see boxed areas in the right-hand side of the displacement graphs in Figures 3D-F). For all three *Plasmodium* species, the percentage of the population moving these large distances was maximal at 5-20 min. While in the rodent *Plasmodium* species, 9.8% (*P. berghei*) and 7.7% (*P. yoelii*) of sporozoites move this far at the 10 min time point, only 3.2% of *P. falciparum* sporozoites reach displacements this large (Figure 3D-F). Interestingly, while for both *P. berghei* and *P. falciparum*, these high-displacing sporozoites are no longer observed at 60 min after inoculation (Figure 3D and 3F), *P. yoelii* sporozoites are still moving larger distances, with 3% of sporozoites moving between 50-200 μm in displacement at 60 and 120 min after inoculation (Figure 3E). This difference in sporozoite behavior between the two rodent *Plasmodium* species is consistent with previous studies which showed that the infectivity of *P. yoelii* sporozoites to rodents is significantly higher than that of *P. berghei* sporozoites (16–19) and is in line with the observation of a persistent exit of *P. yoelii* sporozoites out of the dermis into the circulation for over 2 hours after inoculation (10).

### High gliding speed of *P. yoelii* sporozoites at late time points

The time-lapse images of *P. berghei*, *P. yoelii* and *P. falciparum* sporozoites moving through the dermis of mice were used for analysis of sporozoite gliding speed. Of note, this present analysis of *P. berghei* sporozoite speed was done with a larger set of time-lapse images than the data set used for the speed analysis of tracks generated by the automated tracking method seen in Figure 2D, which was performed on a smaller data set, to match the manual analysis done previously (3). As Figure 4 shows, at 5-10 min after inoculation, sporozoites of all three *Plasmodium* species move at approximately 1.5 μm/s. For *P. berghei*, this speed is stable over the first 30 min and drops off to below 1 μm/s at 60-120 min after inoculation (Figure 4A), similarly to what was described previously (3). In contrast, *P. yoelii* and *P. falciparum* gliding speeds stay more constant over time (Figure 4B,C), as also shown by linear regression of the apparent speed (Figure 4D). At the 120 min time point *P. yoelii* moves significantly faster than *P. berghei* and *P. falciparum* sporozoites (Figure 4A-C), again consistent with a persistent exit of *P. yoelii* sporozoites out of the dermis for over 2 hours after sporozoite inoculation (10).

**Figure 4.**
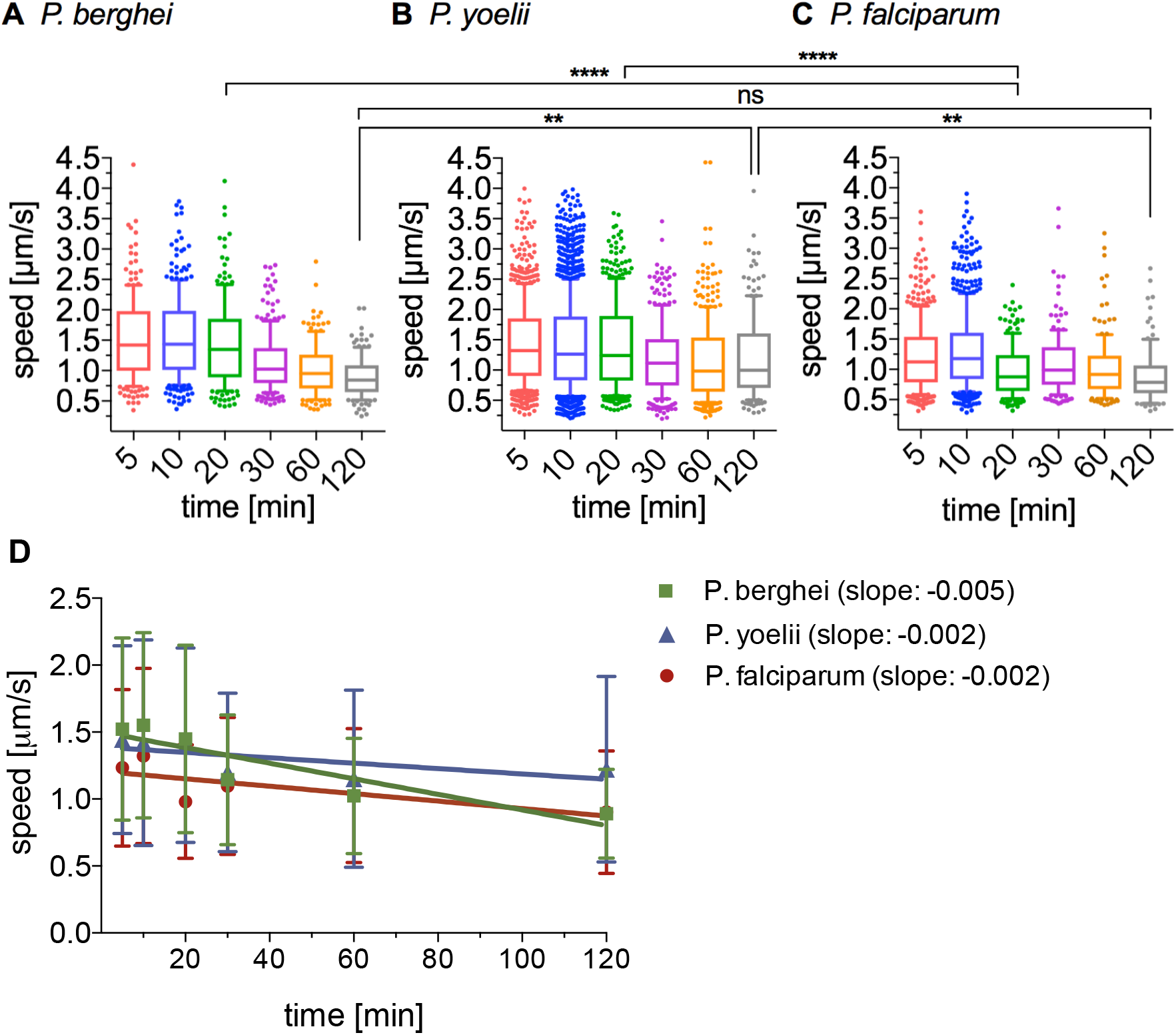
Apparent speed of *P. berghei*, *P. yoelii* and *P. falciparum* sporozoites moving in the dermis 5 to 120 min after intradermal inoculation. **(A-C)** Speed of *P. berghei* (A), *P. yoelii* (B) and *P. falciparum* (C) sporozoites 5-120 min after inoculation in box and whisker plots displaying the 10-90 percentile and individual outliers, with horizontal line showing median displacement at each time point. Comparisons among the species showed statistically significant difference in sporozoite speeds at the 20 min time point (Kruskal-Wallis test, p<0.0001 for both comparisons) and the 120 min time point (Kruskal-Wallis test, p<0.006 for both comparisons). **(D)** Linear regression of apparent speed shows a more rapid decrease for *P. berghei* (slope = −0.005) than for *P. yoelii* and *P. falciparum* (slopes = −0.002). A varying number of videos were processed for each time point after inoculation, using the same data set used for Figure 3: **(A)** 5 min (3 videos/203 tracks), 10 min (6 videos/292 tracks), 20 min (4 videos/184 tracks), 30 min (6 videos/235 tracks), 60 min (7 videos/162 tracks), and 120 min (6 videos/151 tracks). **(B)** 5 min (7 videos/714 tracks), 10 min (16 videos/1514 tracks), 20 min (4 videos/415 tracks), 30 min (4 videos/319 tracks), 60 min (4 videos/255 tracks), and 120 min (4 videos/180 tracks). **(C)** 5 min (6 videos/389 tracks), 10 min (6 videos/464 tracks), 20 min (3 videos/184 tracks), 30 min (2 videos/195 tracks), 60 min (2 videos/239 tracks), and 120 min (2 videos/103 tracks)

### Motility of *P. berghei*, *P. yoelii* and *P. falciparum* sporozoites

As we have described previously, the frequency of motile of *P. berghei* sporozoite motility decreases over time after their inoculation into the dermis (3). We investigated whether *P. yoelii* and *P. falciparum* show the same trend in their motility. Videos of time courses were manually counted to determine the number of sporozoites that are motile and non-motile. The data presented include complete time course experiments spanning 5 min to over 120 min after intradermal injection of sporozoites (Figure 5). To illustrate the percentage of sporozoites leaving the field of view, the data are shown as percentages of the absolute number of sporozoites observed at 5 min after inoculation. We found that compared to *P. berghei*, which, over the course of two hours, lose the ability to move in the dermis, the percentage of motile *P. yoelii* and *P. falciparum* does not rapidly drop off and remains similar at later time points, with 30% to 20% of sporozoites moving at 60 to 120 min after inoculation, respectively. This is consistent with the observation of a slow trickle of *P. yoelii* sporozoites out of the dermis over more than 3 hours after inoculation (10) and suggests that the observed higher infectivity of *P. yoelii* sporozoites compared to *P. berghei* may reflect their biology in the skin in addition to their increased infectivity in the liver (10,16–19).

**Figure 5.**
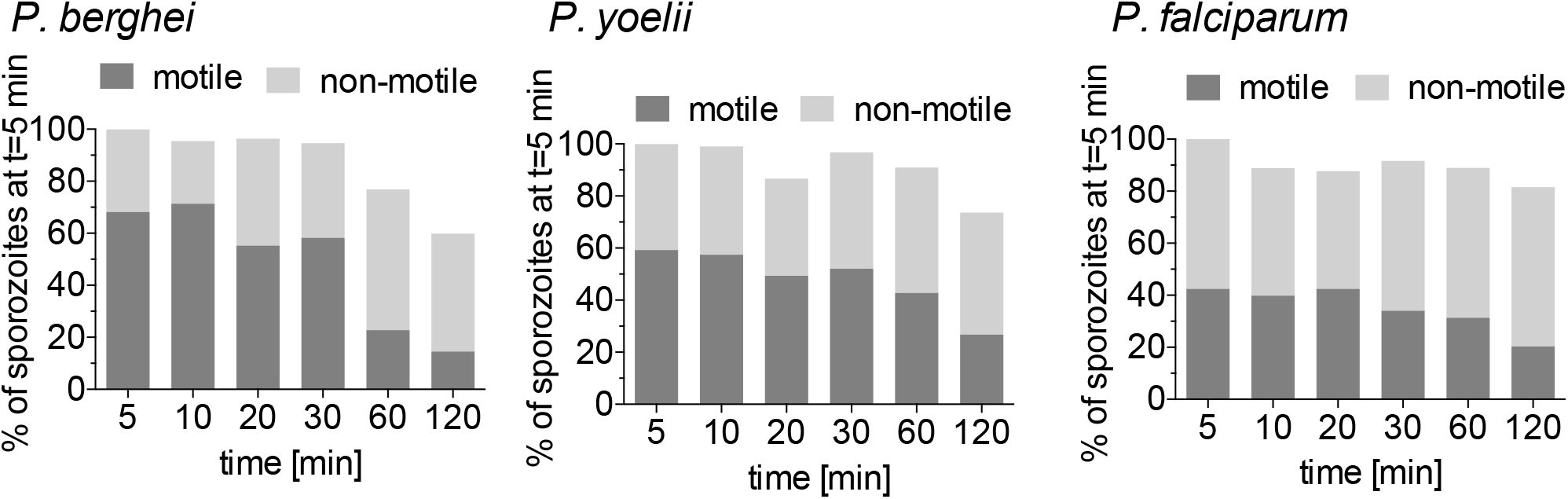
Motility Frequency Over Time of *P. berghei*, *P. yoelii* and *P. falciparum* Sporozoites. The number of motile and non-motile sporozoites moving in the dermis of mice was manually counted in timecourses spanning 120 minutes and is displayed as percentage of total sporozoites observed 5 min after inoculation, for *P. berghei* (left panel), *P. yoelii* (center panel), and *P. falciparum* (right panel). The data shown only includes complete imaging sessions over 120 min after intradermal injection of sporozoites. (*P. berghei*: n=2, *P. yoelii* n=4, *P. falciparum* n=2).

### Blood and lymphatic vessel entry by *P. berghei*, *P. yoelii* and *P. falciparum*

Our previous work had shown that *P. berghei* sporozoites interact with CD31+ blood vessels in the dermis and that while sporozoites are in the vicinity of blood vessels, their motility is more constrained and more circular, likely to optimize contact with dermal capillaries (3). To investigate whether *P. yoelii* and *P. falciparum* sporozoites interact with dermal blood vessels in mouse skin, we imaged these sporozoites in conjunction with fluorescently labeled dermal vascular endothelia. We found that similar to *P. berghei*, both *P. yoelii* and *P. falciparum* engage with CD31 blood vessels by frequently circling around the vessel (Figure 6A and Videos S4-S6). We further quantified entry into blood and lymphatic vessels by the sporozoites, classifying these events similarly to what was previously described, with blood vessel entry being defined by a sudden increase in speed and disappearance of the sporozoite out of the field of view and lymphatic entry being defined by the switch from directed forward movement to sideward drifting of the sporozoite at a low velocity (3,4) (see Videos S7-S8 for time-lapse videos of *P. yoelii* and *P. falciparum* blood vessel invasion events; *P. berghei* blood vessel invasion events can be seen in (3). These events were quantified as a percentage of motile sporozoites in 4 min videos, and data was pooled from videos acquired at 5 min, 10 min and 20 min after intradermal inoculation, thus covering a total imaging time of 12 min as described previously (3). On average 3.1 % and 3.0 % of *P. berghei* sporozoites exit through the blood vessel and lymphatics, respectively during a 4 min video recorded at 10 min post-sporozoite inoculation (Figure 6B, left panel). Both *P. yoelii* and *P. falciparum* sporozoites were found to show similar exit rates (Figure 6B, left panel). We found that 2.6 % of *P. yoelii* sporozoites exit through the blood vessel, while *P. falciparum* sporozoites were found to enter blood vessel at a slightly higher rate than the rodent parasites, with 5.5 % of *P. falciparum* sporozoites entering blood vessels during the acquired imaging time. Both *P. yoelii* and *P. falciparum* sporozoites showed a lower rate of lymphatic invasion (1.5 % and 1.7 %, respectively) compared to *P. berghei*. There were no statistically significant differences in the rate of blood or lymphatic vessel entry among the three *Plasmodium* species.

**Figure 6.**
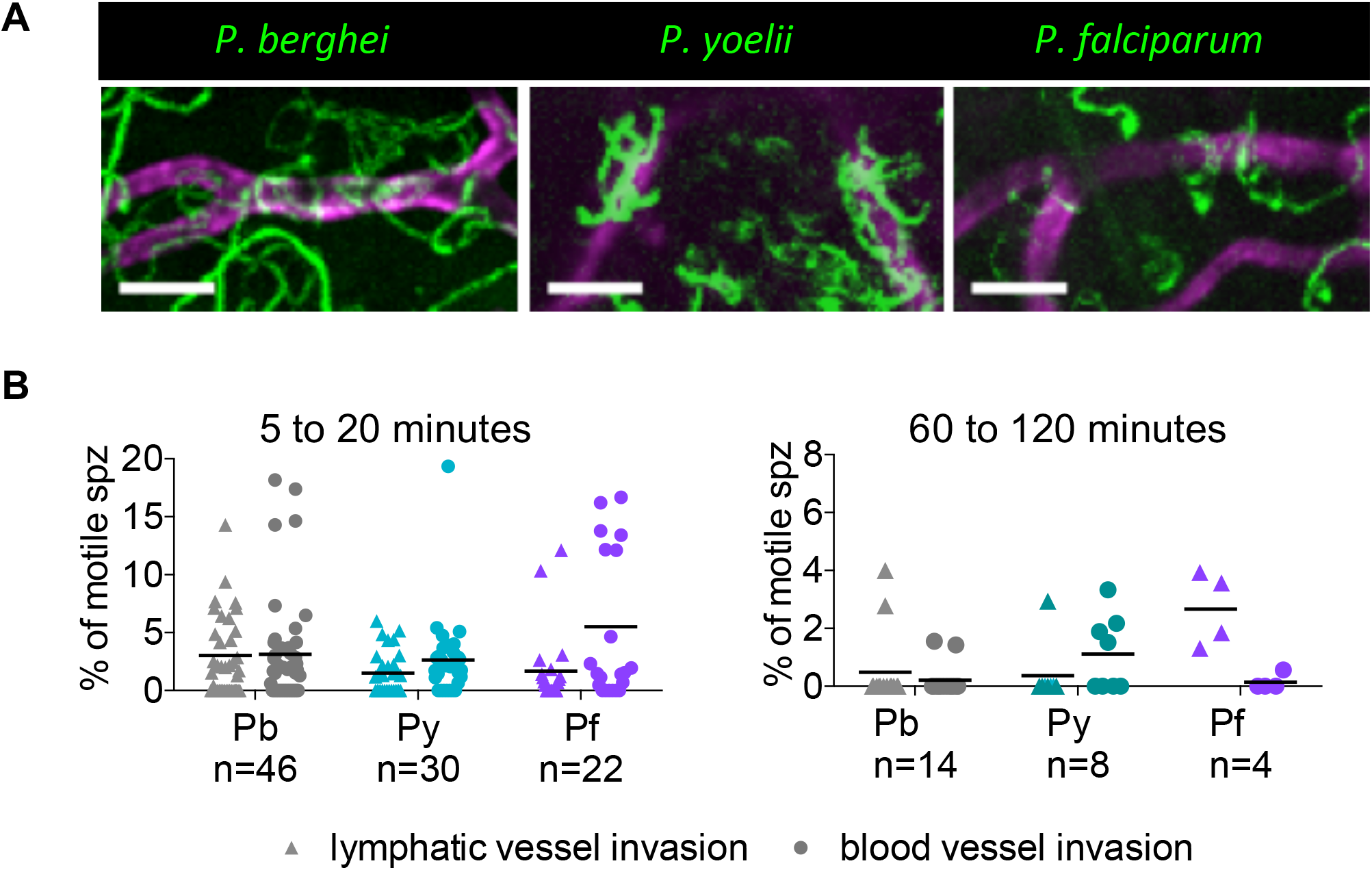
*P. yoelii* and *P. falciparum* sporozoites engage with and enter blood vessels in skin. **(A)** Maximum intensity projection shows trajectories of moving *P. berghei*, *P. yoelii* and *P. falciparum* sporozoites (green), engaging with CD31-labeled vascular endothelia (far red). Scale bar, 25 μm. See videos S4-S6. **(B)** Lymphatic and blood vessel invasion of *P. berghei*, *P. yoelii* and *P. falciparum* sporozoites moving in mouse dermis. Combined invasion events observed in 4 min videos recorded at 5, 10 and 20 min after intradermal injection of sporozoites (left panel) and 60 and 120 min after intradermal injection (right panel). n = number of videos scored for each *Plasmodium* species. To view a blood vessel entry event by *P. yoelii* or *P. falciparum*, see videos S7-8. There were no statistically significant differences in lymphatic or blood vessel entry among the three *Plasmodium* species at 5 to 20 min after sporozoite inoculation whereas there was a statistically significant increase in lymphatic entry by *P. falciparum* sporozoites compared to *P. berghei* and *P. yoeli*i sporozoites at 60 to 120 min after inoculation (p<0.05) (Kruskal-Wallis test).

Given that *P. yoelii* showed more constant gliding speed throughout the first two hours after inoculation and a higher number of sporozoites displacing large distances at 60 to 120 min after inoculation, we decided to investigate whether this translates to blood vessel entry at these later time points. Interestingly, while for *P. berghei*, the rate of blood vessel invasion at these later time points is significantly reduced compared to 5 to 20 min after inoculation (Kruskal-Wallis test, p=0.0301), this was not the case for *P. yoelii*, where even at 60 and 120 min after inoculation sporozoites were still invading blood vessels (Figure 6B, right panel), consistent with the observed higher infectivity of *P. yoelii* sporozoites to rodents compared to *P. berghei* (16–19)

### *P. falciparum* sporozoite motility in the dermis of a humanized mouse model

Thus far our investigations demonstrate that *P. falciparum* sporozoites move and enter blood vessels in mouse skin. To more thoroughly investigate the nature of *P. falciparum* sporozoite motility in vivo, we developed an *in vivo* humanized mouse model of *P. falciparum* skin infection. This was accomplished by grafting human neonatal foreskin onto immunocompromised NOD/scid gamma (NSG) mice, which can readily accept human skin grafts (20). Importantly, the grafted human skin retains the human vasculature and the blood supply to the graft is restored through spontaneous anastomosis of murine and human microvessels (21,22). At 4 weeks after grafting the human skin onto the NSG mice, fluorescent labeling of the human vasculature *in vivo* by intravenous injection of anti-human CD31 (which specifically recognizes human endothelial cells) confirmed that many of the blood vessels in the grafted human skin were of human origin (Figure 7A).

**Figure 7.**
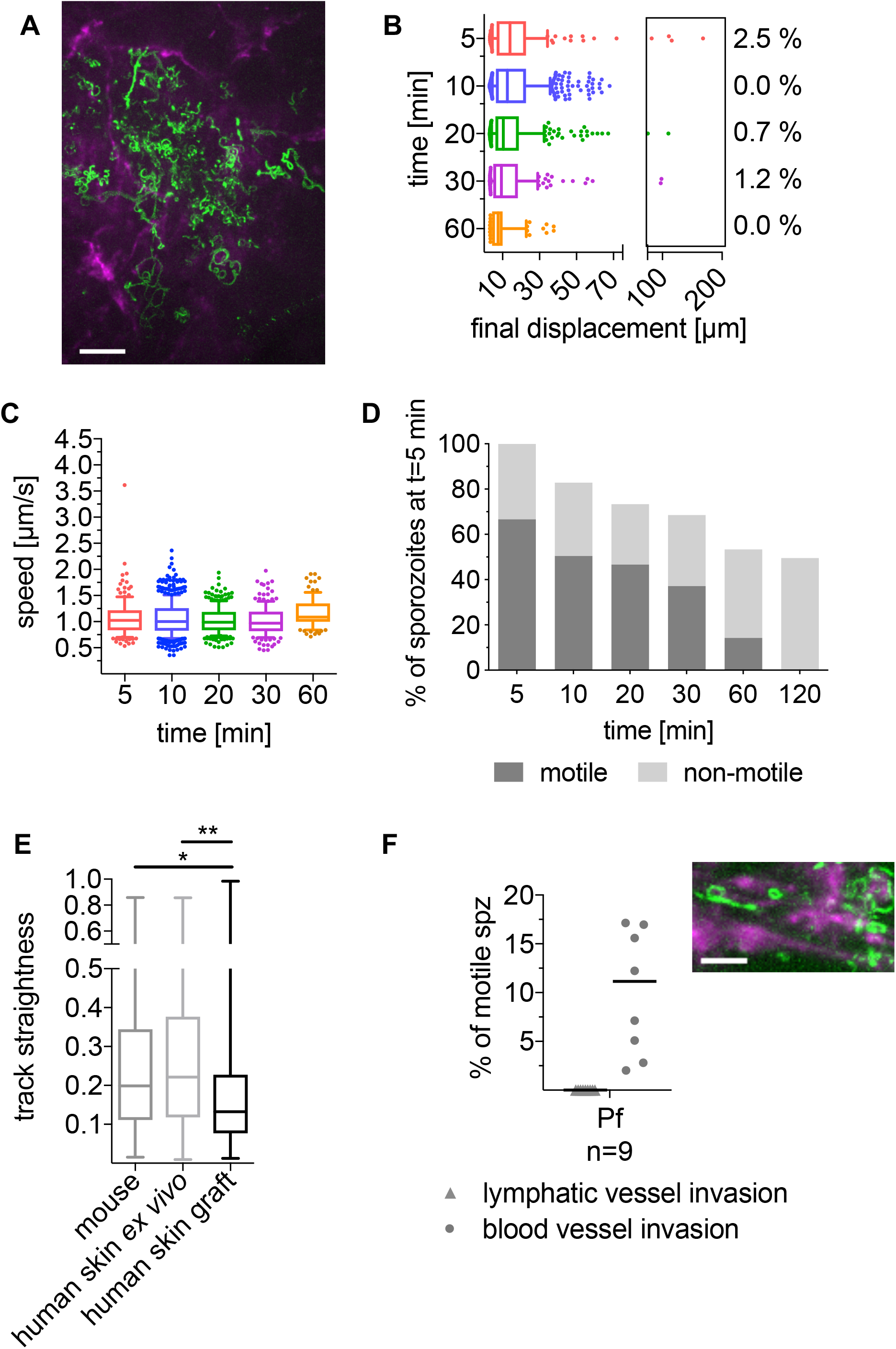
Time-lapse microscopy of *P. falciparum* sporozoites moving in human dermis grafted on NSG mice. **(A)** Maximum intensity projection of video acquired 10 min after inoculation of *P. falciparum* sporozoites visualizes trajectories of parasites, engaging with CD31-labeled vascular endothelia (far red). Scale bar, 50 μm. See video S9. **(B)** Displacement of sporozoites 5 to 60 min after inoculation. Data is displayed in box and whisker plots with the box spanning the 10 to 90 percentile values, the whiskers extending to the highest and lowest observations and the horizontal line showing median. Percentage values indicate the fraction of tracks displacing over 75 microns. **(C)** Apparent speed of sporozoites 5 to 60 min after inoculation. Data is displayed in box and whisker plots with whiskers showing 10 to 90 percentiles and horizontal line showing the median speed. For panels **B**–**C**, data shown originates from the analysis of 3-4 videos per time point and a varying number of tracks were processed for each time point after inoculation: 5 min (157 tracks), 10 min (568 tracks), 20 min (286 tracks), 30 min (171 tracks), 60 min (101 tracks). No motile sporozoites were observed 120 min after inoculation. **(D)** Proportion of motile and non-motile sporozoites was manually counted and is displayed as percentage of sporozoites observed 5 min after inoculation. The data shown was obtained from one complete imaging session over 120 min after intradermal injection of sporozoites. See video S9. **(E)** Sporozoite track straightness, the ratio of displacement to track length of *P. falciparum* sporozoites moving in mouse dermis, human skin *in vitro*, and human skin graft, at 5-10 min post intradermal injection. Data is displayed in box and whisker plots with horizontal line showing the median. Statistical analysis: Tukey’s multiple comparisons test. A varying number of tracks were processed for each environment: mouse skin: 17 videos, 822 tracks; human skin *in vitro*: 12 videos, 761 tracks; human skin graft: 9 videos, 718 tracks. **(F)** *P. falciparum* sporozoites engage with and enter blood vessels in human skin graft. Maximum intensity projection shows trajectories of moving *P. falciparum* sporozoites (green), engaging with human CD31-labeled vascular endothelia (far red). Scale bar, 25 μm. See video S10. Graph shows lymphatic and blood vessel entry events of *P. falciparum* sporozoites in human skin graft. Combined invasion events observed at 5-20 min after intradermal injection in n number of videos. To view a blood vessel invasion event *P. falciparum* in human skin graft, see video S11.

*P. falciparum* sporozoites were injected intradermally into the grafted human skin and intravital imaging performed as previously described (21). *P. falciparum* sporozoites were motile and moved through the grafted human skin tissue. Nonetheless, unlike in the rodent dermis, where at 5 min and 10 min after inoculation the mean displacement of the *P. falciparum* sporozoite population was 22.5 μm and 25.33 μm, respectively; in the human skin graft, the mean displacement at these same time points were 18.6 μm and 16.6 μm, respectively. Indeed, comparison of *P. falciparum* sporozoite displacement in mouse tissue versus the human xenograft was significantly lower in the xenograft at all corresponding time points (compare Figures 7B & 3F: Kruskal-Wallis comparisons: 5 min: *p*<0.05; 10 min: p<0.0001; 20 min: p<0001; 30 min: p<0.005; 60 min: p<0.0001). Additionally, *P. falciparum* sporozoites moved with significantly reduced speed at most time points in human skin compared with what was observed in mouse skin (compare Figures 7C & 4C: Kruskal-Wallis comparisons: 5 min: *p*<0.002; 10 min: p<0.0001; 20 min: p<0.0001; 30 min: ns; 60 min: p<0.02).

The reduced displacement prompted us to analyze the track straightness in order to determine whether these sporozoites were moving in more constrained paths or simply not moving large distances, both of which would result in decreased displacements. We hypothesized that in the human skin grafts, the anastomoses between human and rodent vessels could result in larger, more tortuous vessels, such that sporozoites in the grafted skin could be in closer proximity to vessels and thus be displaying the more constrained motility we had previously observed when *P. berghei* sporozoites were near vessels (3). We therefore measured track straightness, the ratio of displacement to track length (23), to quantify the confinement of motility in grafted human skin and mouse skin. If sporozoite tracks are more constrained, the ratio will be smaller, while straight sporozoite trajectories would result in a larger ratio. This analysis showed that indeed *P. falciparum* sporozoite track straightness in the human skin graft is significantly reduced, compared with the track straightness in mouse skin (Figure 7E), indicating that *P. falciparum* sporozoites move in more constrained circular paths in the grafted human skin.

To determine whether this was due to the human skin environment or to the blood vessels that form in the graft, experiments were performed on human skin specimen *ex vivo* prior to grafting onto mice (Figure S4). Though these specimens do not have a functioning blood supply, the tissue remains alive, being sustained by the oxygen and glucose in the surrounding medium. Interestingly, in the human skin *ex vivo*, *P. falciparum* sporozoite displacement was similar to that in mouse skin and significantly higher than that observed in the human skin xenograft (compare Figures 7B & S4C: Kruskal-Wallis comparisons: 5 min *p*<0.05; 10 min p<0.0001; 20 min p<0001; 30 min p<0.0001; 60 min p<0.0001). We also compared track straightness of sporozoite trajectories in human skin ex vivo to grafted human skin. Interestingly, sporozoite tracks in human skin *ex vivo* were significantly less confined than those in the grafted human skin, similar to what we observed in mouse skin (Figure 7E). Thus, the lower average displacement of *P. falciparum* sporozoites in human skin xenografts appears to be specific to grafted human skin that possess a blood supply.

We next quantified the number of motile sporozoites over time, manually counting motile and non-motile sporozoites and expressing these data as a percentage of the total sporozoites observed at the 5 min time point. Interestingly, over 60 % of inoculated *P. falciparum* sporozoites were motile in the grafted human dermis (Figure 7D), compared with the 40 % *P. falciparum* motile sporozoites in rodent dermis (Figure 5). Moreover, in the grafted human skin, the number of *P. falciparum* sporozoites leaving the field of view was increased compared with rodent dermis, as only ~50% of sporozoites remained at 120 minutes after inoculation. This is likely due to the higher number of motile sporozoites and the larger percentage of *P. falciparum* sporozoites invading blood vessels. As shown in Figure 7F, more than 10 % of motile *P. falciparum* sporozoites invade blood vessels in the human skin graft compared with 5.5% of motile *P.falciparum* in mouse skin. Note that there was no lymphatic invasion in the skin graft since lymphatic vessels are not reconstituted after surgery (24).

## Discussion

Sporozoite motility is essential for successful localization to and entry into blood vessels at the inoculation site. To date, studies of this phase of infection have focused on the more tractable rodent parasite *P. berghei*. Here we extend these studies to the human malaria parasite *P. falciparum* and to *P. yoelii*, a rodent parasite that is highly infectious to the laboratory mouse. Our comparative analysis enabled the identification of components of sporozoite motility that are conserved across species, as well as behaviors that are species-specific or specific to a particular host-parasite combination. Overall, we find that in contrast to the liver and blood stages of malaria, the skin is not a species-specific barrier to infection, with our demonstration of normal motility and blood vessel invasion of *P. falciparum* sporozoites in mouse skin. Thus, the first observations of *P. falciparum in vivo* described in this study likely recapitulate the initial phase of malaria infection in humans.

Many components of sporozoite motility described for the rodent species were observed with *P. falciparum* sporozoites moving in rodent dermis. Sporozoite gliding speed and displacement of *P. falciparum* was similar to that of the rodent parasites at all time points after inoculation. Although the percentage of *P. falciparum* sporozoites reaching displacements larger than 50 μm was lower than the respective value for *P. berghei* and *P. yoelii* at each time point, this trend did not reach statistical significance. For all three *Plasmodium* species, maximal displacement was seen 10 min after sporozoite inoculation, suggesting that sporozoites undergo a phase of activation in the dermis. Although the causal factors remain unknown, it may be linked to the finding that exposure to albumin activates sporozoite motility (25). Importantly, *P. falciparum* sporozoites invade blood and lymphatic vessels in mouse skin at a rate comparable to the rodent parasites and were seen to interact with blood vessels in a similar fashion. These data suggest that sporozoites recognize blood vessels, however, precisely what sporozoites are sensing is not known. Nonetheless, the robust entry of *P. falciparum* sporozoites into rodent blood vessels suggests it is something common to mice and primates. The finding that *P. falciparum* sporozoites in mouse skin recapitulate, at least to some degree, canonical sporozoite behavior at the inoculation site, is consistent with a recent study demonstrating *P. falciparum* sporozoites delivered by mosquito bite were infectious in mice with humanized livers (26). However, *P. falciparum* sporozoites have a somewhat decreased infectivity in this model, which could be do to incomplete engraftment of human hepatocytes or to abnormal behavior of sporozoites in mouse skin. Our study suggests that this decreased infectivity is due to incomplete engraftment of human hepatocytes and overall, our data suggest that the way sporozoites move in the skin and recognize blood vessels are conserved features for *Plasmodium* parasites.

Though the behavior of human and rodent malaria sporozoites in mouse skin share many features, we also observed differences that were host-specific. For both rodent parasites, the proportion of the inoculum that is motile within 5 to 10 minutes of inoculation was approximately 70 % and 55 % for *P. berghei* and *P. yoelii* sporozoites, respectively. In contrast, approximately 40% of *P. falciparum* sporozoites were motile shortly after their inoculation and this percentage remained fairly consistent throughout the 60 minutes after sporozoite inoculation. The continued motility of *P. falciparum* sporozoites over 60 minutes is similar to *P. yoelii*, whereas the proportion of *P. berghei* sporozoites, that continue to be motile over a 60-minute timeframe, significantly decreases. Nonetheless, the overall lower percentage of motile *P. falciparum* sporozoites suggests that there may be some host-specific signals that enhance motility after inoculation into the dermis. Interestingly, when inoculated into the grafted human skin these differences in *P. falciparum* sporozoite motility disappear: A higher proportion of the inoculum is motile, with over 60 % of sporozoites motile at 5 min after inoculation, suggesting that there are indeed host-specific components to sporozoite motility in the skin. Nonetheless, we found that there are some limitations to the humanized mouse model. Foremost are the lower displacements and speed of *P. falciparum* sporozoites observed in the human skin graft. We think this is due to the anastomoses between human and rodent vessels, which results in larger and more tortuous vessels. Given that these vessels are larger, it is likely that sporozoites in the grafted human skin were in closer proximity to vessels, and this might explain the more constrained motility and lower speed of the *P. falciparum* sporozoites in human skin compared with the mouse skin. Interestingly, speed and displacement of *P. falciparum* in human skin *ex vivo* were similar to what we observed in mouse skin, further supporting this hypothesis. Thus, it is likely the anatomy of the grafted and anastomosed vessels in the human skin graft that is responsible for the decreased displacement and speed of sporozoites in this setting.

Comparison between the two rodent malaria species, both of which are commonly used for vaccine studies, demonstrated that *P. yoelii* sporozoites are able to move more persistently, with higher speed and larger displacements at later time points. We also observed more blood vessel invasion events by *P. yoelii* sporozoites at later time points. These data suggest that *P. yoelii* sporozoites have a longer period of time during which they are infectious post-inoculation. While previous studies using intravenously inoculated sporozoites demonstrated that *P. yoelii* sporozoites develop better in the liver compared to *P. berghei* (16–19), our results suggest that the higher infectivity of *P. yoelii* sporozoites is also due to their behavior in the skin, further supporting the notion that sporozoite motility in the dermis is a valid correlate for infectivity of the parasite. Recently a number of transgenic rodent parasite lines expressing *P. falciparum or P. vivax* CSP have been developed for pre-clinical vaccine testing in the rodent model (27–29). The different biology of the two rodent malaria species could impact the interpretation of antibody-based interventions performed in the rodent model and should be considered when evaluating vaccine candidates targeting sporozoites. Given the results of our comparative analysis, a *P. yoelii* parasite expressing *P. falciparum* proteins such as CSP (30) may be the more robust model.

The quest for a potent malaria vaccine remains a global priority to decrease clinical disease and mortality and enable elimination. Despite this, the development of a highly effective and durable malaria vaccine has proven challenging. In part, this goal has been hampered by the lack of robust pre-clinical models for vaccine testing. Recent efforts have focused on characterizing the parasite-specific antibody response in vaccinated or naturally exposed individuals (reviewed in (31,32)), which has opened up new possibilities for epitope-focused vaccine design and passive antibody therapies, but there is a scarcity of models to test protective efficacy prior to clinical trials. Here we describe an *in vivo* assay in mice that can be used to screen human monoclonal antibodies and vaccine candidates with *P. falciparum* sporozoites. This is a significant step forward and has several advantages over currently used assays for pre-clinical testing of inhibitory activity on *P. falciparum* sporozoites. *In vitro* assays quantifying cell traversal and invasion suffer from the low infectivity of *P. falciparum* sporozoites *in vitro* and our lack of knowledge on how to activate *P. falciparum* sporozoites for these processes once they have been dissected from mosquito salivary glands. Significant progress was made with the development of a humanized mouse model (33), however, the mice used for these assays are expensive and immune-compromised, making it difficult to evaluate vaccine candidates using this model. Our model, using immune-competent mice, enables the use of large numbers of mice due to their low-cost relative to the humanized mice, and evaluation of vaccine candidates due to their fully competent immune system. Moreover, the automated sporozoite tracking tool we developed, facilitates more rapid data analysis. Together this makes for a platform that will enable higher throughput testing of vaccines and antibodies targeting human malaria sporozoites, prior to costly primate or human studies.

## Materials and Methods

All animal experiments were approved by the Johns Hopkins Animal Care and Use Committee.

### Generation of fluorescent *P. yoelii* -mCherry

The targeting plasmid pL1849-mCherry_CSP_, containing sequences that would target the *mcherry* transgene to the *P. yoelii 230p* locus, and the *mcherry* coding sequence flanked by the *P. berghei csp* 5’UTR and the *P. berghei dhfr* 3’UTR was generated as follows: the *5’pbcsp-mCherry-3ʹpbdhfr* cassette was amplified from plasmid pL0047 (RMgm database http://www.pberghei.eu), with primers pL0047-PbCS-mCherry-F and pL0047-PbCS-mCherry-R (see Table S1 for primer sequences), which added XmaI sites for cloning into pL1849 (34), which contained the *P. yoelii* 230p locus (PY03857) 5’− and 3’-targeting sequences (Figure S1). The resulting construct pL1849-mCherry_CSP_ was released from the plasmid through digestion with BstXI and ScaI and used for transfection into the *P. yoelii* line Py17X_GIMO_ (line 1923cl1) (34), according to standard procedures (35). The following day, negative selection with 1 mg/ml 5-FC in drinking water (36) was used to enrich for parasites that integrated the *PbCSP-mCherry-3ʹpbdhfr* cassette and excised the *hdhfr*::*yfcu* locus through double cross-over homologous recombination. The resulting PymCherry line was cloned and diagnostic PCRs confirmed integration and absence of the h*dhfr*::y*fcu* marker (Figure S1). See Table S1 for all primer sequences. Clone D2 was used for all experiments in the present study.

### Intravital imaging

Sporozoites inoculated into the ear pinna of mice were imaged as described previously (3,37). Briefly, anesthesia of 4− to 5-week-old C57BL/6 mice from Taconic was done by intraperitoneal injection of ketamine/xylazine (35–60 μg ketamine/gram body weight), and sporozoites were injected intradermally into the ear pinna in a total volume of 0.2 μl using a NanoFil-10 μl syringe with a NF33BV-2 needle (World Precision Instruments). All sporozoite injections were completed within 30 min of salivary gland dissection to ensure that imaging and parasite conditions remained constant. The ear pinna was taped to a coverslip and the mouse was mounted on the platform of an inverted Zeiss Axio Observer Z1 microscope with a Yokogawa CSU22 spinning disk and a preheated temperature- controlled chamber at 30 °C. Parasites were imaged with a 10× objective and magnified using a 1.6 Optovar, resulting in an x and y imaging volume of 500 μm × 500 μm. Stacks of 3 slices spanning a total depth of 30–50 μm were imaged. Exposure time per slice was 100 to 150 ms and stacks were captured at approximately 1 Hz over a total time of 4 min using an EMCCD camera (Photometrics, Tucson, AZ, United States) and 3i slidebook 5.0 software. Indicated time points after inoculation refer to the starting time of the 4-min acquisition. In order to visualize dermal blood vessels, mice were intravenously injected with 15 μg of rat anti-mouse CD31 coupled to Alexa-Fluor-647 (clone 390, Biolegend, San Diego, CA, United States) approximately 2 hr prior to imaging.

### Human skin graft

Each 6-12 week-old female NOD-SCID gamma (NSG) mouse (NOD.Cg-Prkdc^scid^ Il2rg^tm1Wjl^/SzJ, Jackson Laboratories) was anesthetized with inhalation isoflurane (2%) and their entire back and bilateral chest walls were shaved with clippers. To make a recipient wound bed for human donor foreskin engraftment, the skin was sterilized with alchohol (70%) and povidone-iodine (Betadine) and survival surgery was performed to excise a 1.5 cm × 1.2 cm rectangular area of full thickness skin below the shoulder blades. Each normal human foreskin specimen was obtained from normally discarded foreskin from a male newborn at the Johns Hopkins Hospital and was placed into RPMI at 4°C for 1 to 4 hours prior to use. Each foreskin specimen was cut into a 1.4 × 1.1 cm rectangular area and the subcutaneous fat and fascia sterilly removed with scissors and forceps. Each foreskin graft was placed onto the wound bed on each recipient NSG mouse and secured with interrupted nylon sutures between the skin graft and adjacent mouse skin and muscle fascia. Topical bacitracin ointment was applied to the human skin graft site and the graft was then covered with a non-adherent dressing (Telfa, Covidien) followed by an adhesive bandage (Band-aid, Johnson & Johnson). Each NSG mouse with engrafted human foreskin was singly housed in an autoclaved cage with with water supplemented with baytril (0.5 mg/ml) and aspartame sweetener. After 2 weeks, bandages and sutures were removed and mice were monitored for another 2 weeks to ensure graft survival prior to using the mice in subsequent experiments.

### Imaging of sporozoites in human skin graft

Anesthesia of NSG mice with human skin graft was done by intraperitoneal injection of ketamine/xylazine (35–60 μg ketamine/gram body weight). Similar to a previously described protocol (21), a midline dorsal incision around three sides of the human skin graft was made without disrupting the lateral dermal blood supply. The skin human graft was separated from the underlying tissue and the microvasculature within the graft was exposed through dissection of the connective tissue adhered to the graft. *P. falciparum* sporozoites were then injected into the dermis in a total volume of 0.2 μl using a NanoFil-10 μl syringe with a NF33BV-2 needle (World Precision Instruments). The human skin was then moistened with RPMI medium, immobilized to a coverslip with two small drops of superglue and the edges of the human skin graft were covered with lubricant Puralube Vet ointment (Dechra, NDC 17033-211-38) to avoid drying out of the tissue. The mouse was mounted on the platform of an inverted Zeiss Axio Observer Z1 microscope with a Yokogawa CSU22 spinning disk and imaging was performed as described above. To visualize human dermal blood vessels, mice were intravenously injected with 15 μg of mouse anti-human CD31 coupled to Alexa-Fluor-647 (clone WM59, Biolegend, San Diego, CA, United States) approximately 2 hrs prior to imaging.

### Image processing

Time-lapse stacks with 3-5 slices spanning a total time of 4 min were projected into a single z-layer and the resulting two-dimensional data set was exported as tiff files. Fiji software, which is freely available (https://fiji.sc), was used to remove background and threshold the images. The thresholded time series was analyzed using ICY 1.8.6.0 software (BioImage Analysis Unit; Institut Pasteur, Unite d’analyse d images quantitative; 25,28 Rue du Docteur Roux 75015 Paris, France) for automated spot detection and track generation. A more detailed protocol of the image processing steps performed and configuration details for spot detection and tracking can be found in Supplementary Material. To remove background noise, tracks of less than 30 seconds in total duration, tracks with a total track length of less than 20 μm, tracks with a total displacement of less than 3 μm and tracks with an average speed higher than 4 μm/s were excluded from the analysis. Speed and displacement data was exported to an Excel file and track projections to a common origin were created to visualize parasite dispersal.

### Displacement and Speed

Total displacement was calculated as the Euclidean distance between the first and the last sporozoite location. Sporozoite speed was calculated as the total track length, divided by the track duration. This measure is sensitive to the frame rate with which the video was acquired (38) and to achieve consistent interval times across the entire data set, for the calculation of apparent speed we utilized only videos acquired at 850-1250 ms per frame.

### Track straightness

The track straightness is defined as the ratio of track displacement to total track length and as a result can range between zero (entirely constrained track) and one (the track is a straight line) (23). To calculate the track straightness of *P. falciparum* sporozoite trajectories, the tracks generated for displacement and speed analysis were used.

## Acknowlegdements

We would like to thank Dr. Godfree Mlambo, Dr. Abhai Tripathi and Chris Kizito, the team of the parasitology and insectary core facilities at the Johns Hopkins Malaria Research Institute for their outstanding work and Bloomberg Family Philanthropies for their support of these facilities. We acknowledge the Johns Hopkins School of Medicine Microscopy Facility (MicFac) and thank Dr. Scott Kuo and Barbara Smith for their invaluable assistance. This work was supported by the National Institutes of Health (R01 AI132359 to PS, R01AR069502 and R01AR073665 to LSM and K01AR073924 to NKA), by a Johns Hopkins Malaria Research Institute fellowship (CSH) and by the Bloomberg Family Philanthropies.

## Supplementary Materials

### Supplementary Methods: Automated sporozoite tracking method

**Fig. S1. Generation and verification of a *Plasmodium yoelii* line expressing mCherry under control of the *P. berghei csp* promoter.**

**Fig. S2. Image processing and automated tracking method.**

**Fig. S3. Sporozoite displacement obtained by tracking of complete 4-min tracks compared to tracking of total sporozoite population.**

**Fig. S4. Motility of *P. falciparum* sporozoites moving in the dermis of human dermis *in vitro.***

**Table S1. Primers used for genotypic analysis of the Py-mCherry line.**

### Supplementary Videos

**Video S1.** Time-lapse microscopy of *P. berghei* sporozoites (green) starting 10 min after intradermal inoculation, together with CD31-labeled vascular endothelia (far red). Scale bar, 50 μm. Maximum projection shown in Figure 1A.

**Video S2.** Time-lapse microscopy of *P. yoelli* sporozoites (green) starting 10 min after intradermal inoculation, together with CD31-labeled vascular endothelia (far red). Scale bar, 50 μm. Maximum projection shown in Figure 1B.

**Video S3.** Time-lapse microscopy of *P. falciparum* sporozoites (green) starting 10 min after intradermal inoculation, together with CD31-labeled vascular endothelia (far red). Scale bar, 50 μm. Maximum projection shown in Figure 1C.

**Video S4.** Time-lapse microscopy of *P. berghei* sporozoites (green), engaging with CD31-labeled vascular endothelia (far red). Scale bar, 25 μm. Maximum projection shown in Figure 6A.

**Video S5.** Time-lapse microscopy of *P. yoelii* sporozoites (green), engaging with CD31-labeled vascular endothelia (far red). Scale bar, 25 μm. Maximum projection shown in Figure 6B.

**Video S6.** Time-lapse microscopy of *P. falciparum* sporozoites (green), engaging with CD31-labeled vascular endothelia (far red). Scale bar, 25 μm. Maximum projection shown in Figure 6C.

**Video S7.** Time-lapse microscopy showing entry into CD31-labeled blood vessels (far red) by *P. yoelii* sporozoites (green). Scale bar, 50 μm.

**Video S8.** Time-lapse microscopy showing entry into CD31-labeled blood vessels (far red) by *P. falciparum* sporozoites (green). Scale bar, 50 μm.

**Video S9**. Time-lapse microscopy of *P. falciparum* sporozoites (green) starting 10 min after intradermal inoculation into human skin graft, together with CD31-labeled human vascular endothelia (far red). Scale bar, 50 μm. Maximum projection shown in Figure 7A

**Video S10**. Time-lapse microscopy of *P. falciparum* sporozoites (green), starting 10 min after inoculation into human skin graft, engaging with CD31-labeled vascular endothelia (far red). Scale bar, 25 μm. Maximum projection shown in Figure 7E.

**Video S11**. Time-lapse microscopy showing *P. falciparum* sporozoites (green) entering both human CD31-labeled vessels (far red) and unlabeled mouse vessels in a human skin graft. Scale bar, 50 μm.

