## Supplementary Material for "Quantitative intravital imaging of *Plasmodium falciparum* sporozoites: A novel platform to test malaria intervention strategies"

### Supplementary Methods: Automated sporozoite tracking method

#### 1. Image processing

Timelapse stacks with 3-5 slices spanning a total depth of 30-50 μm were captured at approximately 1 Hz over a total time of 4 min using 3i slidebook 6.0 software (Intelligent Imaging Innovations) and exported as tiff files. Note was taken of the pixel size (pixel size is dependent on camera resolution, using a Photometrics Evolve camera and 16x magnification, the pixel size was 1μm/pixel) and the time interval, i.e. the time elapsed between two captured Z stacks. Using Fiji software, which is freely available https://fiji.sc, the exported raw data as tiff file was opened (File > Import > Image Sequence) and the image was projected into a single Z-layer using a method called maximum-intensity projection (Image > Stacks > Z project > Max Intensity). For background subtraction, an average projection over the Z-dimension was generated (Image > Stacks > Z project > Average Intensity). The resulting average was subtracted from each image of the maximum projected timelapse series (Process > Image Calculator > Subtract). An example is shown in Figure S2 panel B. The threshold of the timelapse series was adjusted (Image > Adjust > Threshold > deselect "calculate threshold for each image"), to remove remaining background signal and an example of a thresholded image is shown in Figure S2 panel C.

#### 2. Spot detection and tracking

The thresholded timeseries was opened using ICY 1.8.6.0 software (BioImage Analysis Unit; Institut Pasteur) and if necessary, the image was converted to a timelapse series (Sequence Operation > Convert to time). Before spot detection was started, pixel size and time interval were set in "Sequence properties". For spot detection, the Spot Detector plugin was used (Detection & Tracking > Spot Tracking > Run the Spot Detector plugin; Detector > "Detect bright spot over dark background" and select scale 3 (~7 pixels) scale 4 (~13 pixels); Output > select "export to SwimmingPool"). An example of an overlay of detected spots over the original thresholded image is shown in Figure S2 panel D. The parameters for sporozoite tracking were: expected false detections per frame: 50; Probability of detection for each particle: 0.9; expected track length: 50; Minimum probability of existence: 0.5; Probability of existence threshold for track termination: 0.0001; Single motion model: expected displacement length in the x-y plane: 10; select "use directed motion"; expected displacement length in the x-y plane: 10; select "re-estimate online"; Expected number of new objects per frame: 5; Expected number of objects in the first frame: 85; Depth of the track trees: 4; Gate factor for association: 4;. These parameters had previously been saved to "tracking con**fig.**xml" and were loaded (Spot Tracking > Interface > advanced Interface > Configuration file > load a configuration file). Spot detection results were used for tracking (Spot Tracking > Detection Source > Select detection results here) and the input of spots created in Spot Detection were chosen. An example of an overlay of created tracks over the original thresholded image is shown in Figure S2 panel E.

#### 3. Track managing and data export

Tracks of less than 30 seconds in duration were selected (TrackManager > Edit > Select track by length > 0-X [X = 30,000/interval time]) and removed (TrackManager > Edit > Delete selection). Furthermore, tracks of less than 20 μm in total track length were removed (TrackManager > add Track Processor > Motion Profiler > select "Filter tracks..." and "Use real units"; Then under "Keep tracks with" select "total displacement > 20" [this removes tracks with a total track length of less

than 20 µm only if the pixel/ µm ratio is 1]). Also, tracks of less than 3 µm in final displacement, were removed (TrackManager > add Track Processor > Motion Profiler > select "Filter tracks..." and "Use real units"; Then under "Keep tracks with" select "net displacement > 3" [this removes tracks with a total track length of less than 20 µm only if the pixel/ µm ratio is 1]).
Speed and displacement data was exported to an Excel file (TrackManager > add Track Processor > Motion Profiler > select "Use real units" > Export statistics) and Track projections to a common origin were created to visualize parasite dispersal file (TrackManager > add Track Processor > Motion Profiler > select "Use real units" > Save graphics in a .png file). An example of a track is shown in Figure S2 panel F. The positional x/y data of all tracks over time was exported (TrackManager > add Track Processor > Track Processsor export track to Excel).

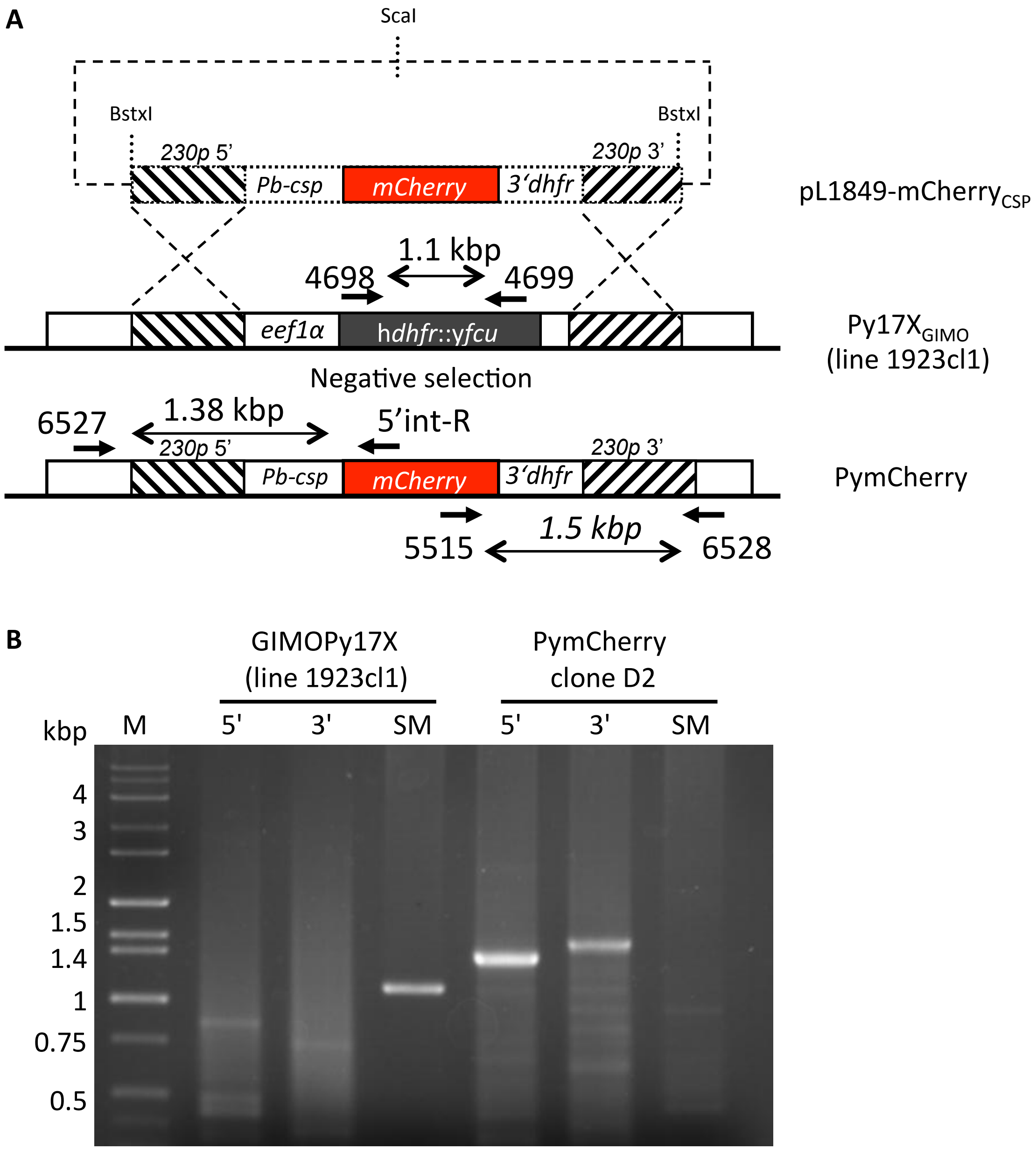

**Figure S1. Generation and verification of a *Plasmodium yoelii* line expressing mCherry under control of the *P. berghei csp* promoter. (A)** Schematic representation of the introduction of an mCherry-expression cassette under the control of the *P. berghei csp*-promoter into the Py17X$_{GIMO}$ (line 1923cl1) *(34)*. Transfection construct pL1849-mCherry$_{CSP}$ containing the *pbcsp-mCherry-3'pbdhfr* cassette was linearized by digestion with BstXI and ScaI and was integrated into the modified *P. yoelii* 230p locus containing the h*dhfr*::y*fcu* selectable marker cassette (grey box) by double cross-over homologous recombination at the target regions (hatched boxes). Negative selection with 5-FC selected for parasites that have the mCherry reporter introduced into the genome and the h*dhfr*::y*fcu* marker removed (PymCherry line). Location of primers used for PCR analysis and sizes of PCR products are shown. **(B)** Following transfection and cloning, diagnostic PCRs confirmed integration as expected in the PymCherry line (clone D2), shown by the absence of the h*dhfr*::y*fcu* marker (amplification of h*dhfr*::y*fcu* with primers 4698/4699) and correct 5′- and 3′-integration PCR product sizes (primer pairs 6527/5'int-R and 5515/6528, respectively). See Table S1 for all primer sequences.

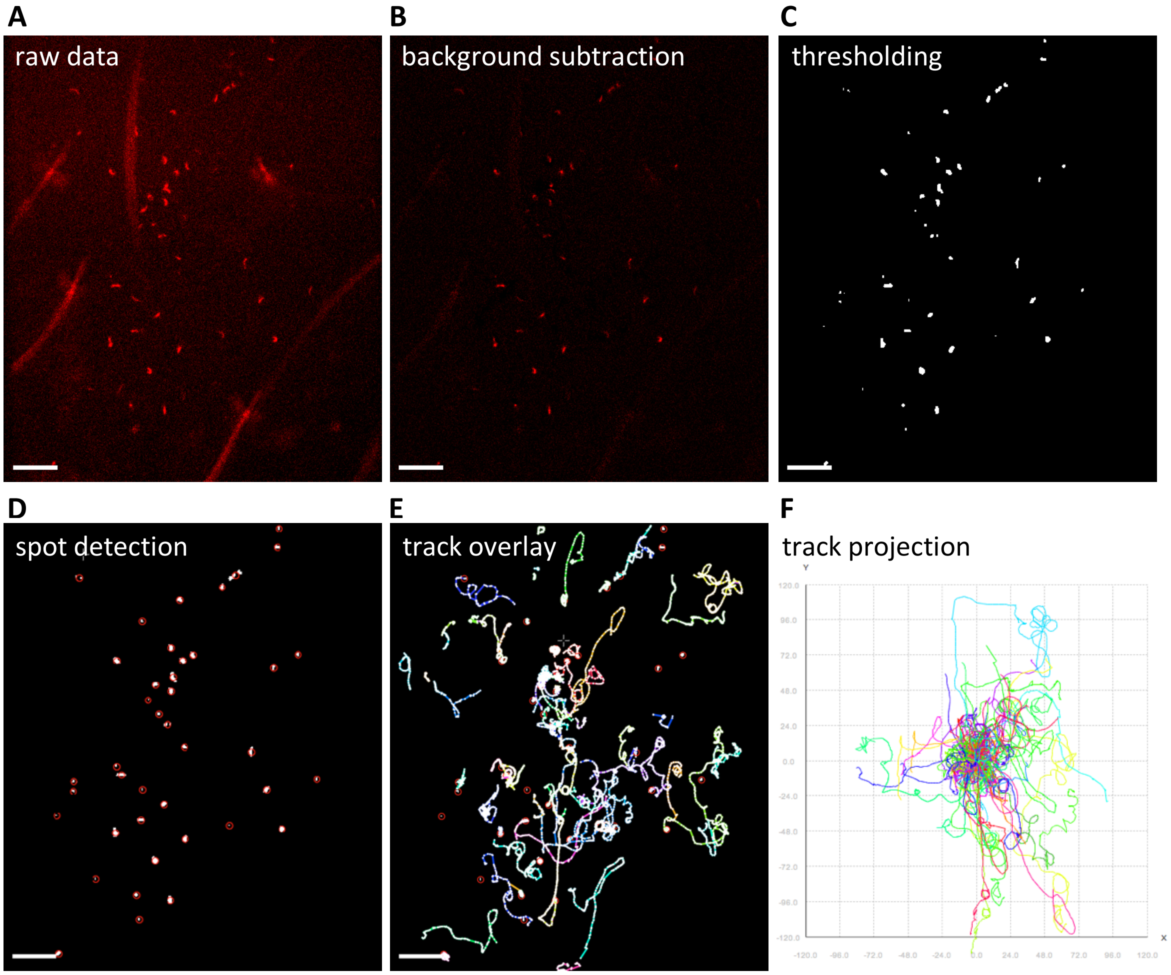

**Figure S2. Image processing and automated tracking method.**
**(A)** First frame of time-lapse microscopy of *P. berghei* sporozoites (red) after intradermal inoculation. After background subtraction **(B)** and thresholding **(C)** using Fiji software, spot detection **(D)** was run using ICY software. **(E)** Tracks were generated using the spot tracking plugin of ICY software. **(F)** Tracks generated were plotted to a common origin, to visualize parasite dispersal. Scale bars, 50 µm

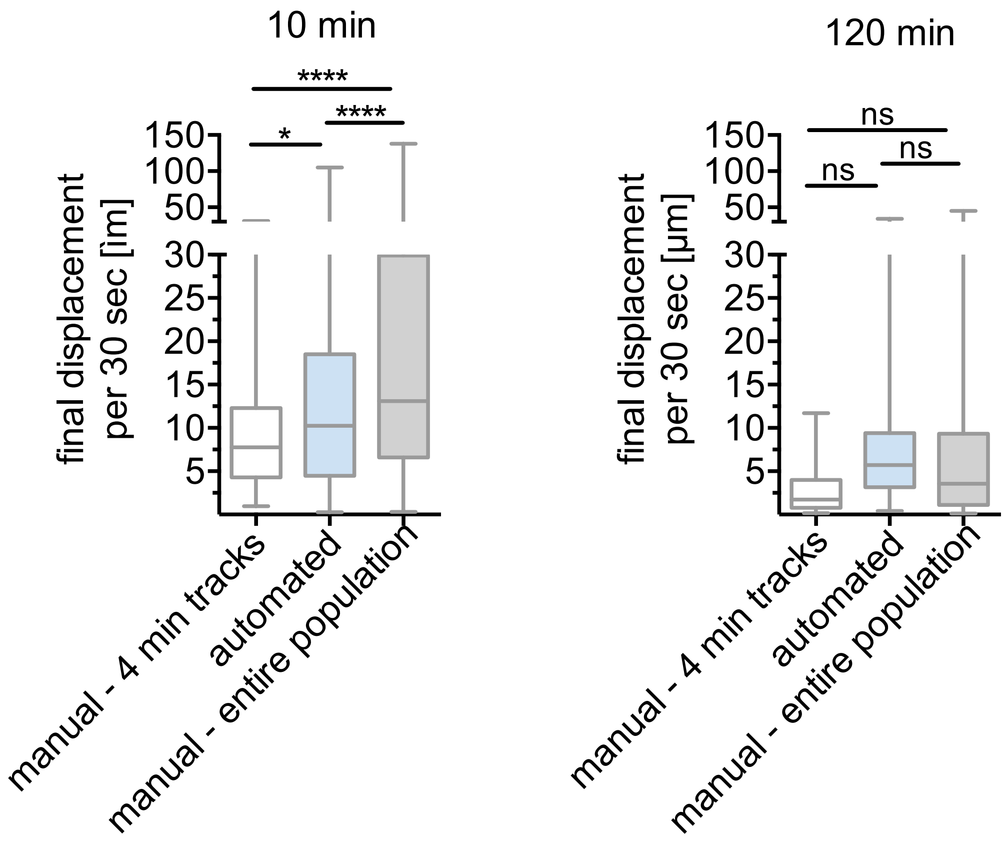

**Figure S3. Sporozoite displacement obtained by tracking of complete 4-min tracks compared to tracking of total sporozoite population.**

Displacement obtained by manual tracking of sporozoites that do not leave the field of view throughout the 4 min video, was compared to automated tracking data, corresponding to the entire sporozoite population and data obtained from manual tracking of the entire sporozoite population. Displacement of tracks **(A)** 10 min and **(B)** 120 min after intradermal inoculation is shown. Data is displayed in box and whisker plots with horizontal line showing median. A varying number of videos were processed for each time point after inoculation: 10 min (6 videos/74 manual-4 min tracks/292 automated tracks/201 manual-entire population), 120 min (5 videos/48 manual-4 min tracks/151 automated tracks/107 manual-entire population). Statistical analysis (Kruskal-Wallis test), * $p<0.05$; **** $p<0.0001$.

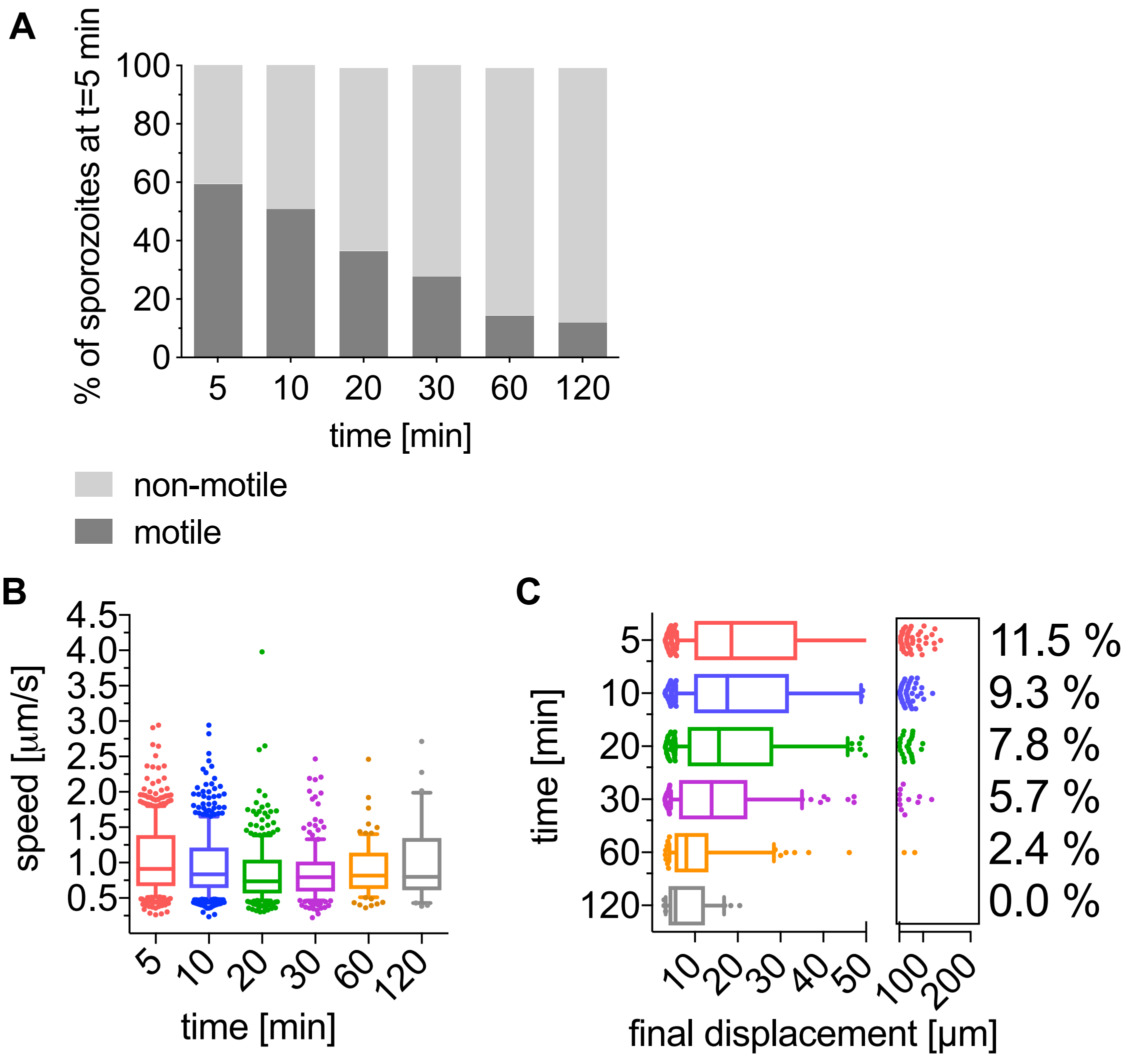

**Figure S4. Motility of *P. falciparum* sporozoites moving in human skin *ex vivo*.** The data shown originates from four complete imaging sessions over 120 min after intradermal injection of sporozoites into *ex vivo* human skin tissue. **(A)** Motile and non-motile sporozoites were manually counted and proportions are displayed as percentage of sporozoites observed 5 min after inoculation. **(B)** Apparent speed of sporozoites 5 to 120 min after inoculation. Data is displayed in box and whisker plots with whiskers showing 10 to 90 percentiles and horizontal line showing the median speed. **(C)** Displacement of sporozoites 5 to 120 min after inoculation. Data is displayed in box and whisker plots with the box spanning the 10 to 90 percentile values, the whiskers extending to the highest and lowest observations and the horizontal line showing median. Percentage values indicate the fraction of tracks displacing over 75 microns.

**Table S1. Primers used for genotypic analysis of the Py-mCherry line.**

| **Generation of Py-mCherry** | |
|---|---|
| **Primer Name** | **Primer Sequence** |
| pL0047-PbCS-mCherry-F | GCACGCCCGGGGCCCTTGCGCCCTTAAGACA |
| pL0047-PbCS-mCherry-R | CGTGCCCCGGGCGAGCTCGGTACCCGAAATTGAA |
| 6527 5'- intgr *py230p*, F | GAAGGATATGAATTAGATCCACC |
| 5'-int-R | ATTGTAAAATTGAGGATGCTTGT |
| 5515 mCherry-F | GCATGGACGAGCTGTACAAG |
| 6528 3'- intgr *py230p*, R | AGACATTGGCATATGAGCAAG |
| 4698 h*dhfr*, F | GTTCGCTAAACTGCATCGTC |
| 4699 y*fcu*, R | GTTTGAGGTAGCAAGTAGACG |
